# First record of the Australian redclaw crayfish *Cherax quadricarinatus* (von Martens 1868) in Hong Kong, China

**DOI:** 10.1101/2020.11.17.387696

**Authors:** YAU Sze-man, LAU Anthony

## Abstract

Invasive freshwater crayfish are spreading rapidly across the world. Here, we report the first record of Australian redclaw crayfish, *Cherax quadricarinatus* (von Martens 1868) in Hong Kong, China. Identification of the captured crayfish was confirmed using external morphological features and molecular analyses. A total of 49 crayfish were captured from a stream pool and a reservoir in Pok Fu Lam Country Park using dip nets and funnel traps. The captured *C. quadricarinatus* ranged from 17.20 mm to 56.40 mm (mean = 30.70) in carapace length and the sex ratio was 1:1. Since this species is globally recognized as an invasive species, a comprehensive survey on its status and invasion front, an investigation into its potential ecological impacts, as well as the formulation of a monitoring and removal strategy, are warranted.

## Introduction

Introduced alien species have been generally recognized as one of the greatest threats to global biodiversity (Dudgeon et al. 2006; Pyšek et al. 2020; Tickner et al. 2020). In addition to interacting with native species through competition and predation, some introduced species can modify habitats and cause community-level shifts in species assemblages and associated ecosystem processes (Sanders et al. 2003; Eastwood et al. 2007).

A wide range of animals and plants have been introduced to Hong Kong, a densely populated metropolitan located on the edge of the tropic. Relatively wild winters and abundant natural habitats enabled the establishment of at least 74 species of invasive species in the city, including mosquito fish *Gambusia affinis*, red imported fire ant *Sloenopsis invicta*, and mile-a-minute weed *Mikania micrantha*. Despite recognizing invasive species as a key threat to the local natural environment, the Hong Kong government does not maintain a detailed database on invasive species and has put forth minimal effort to manage these species (GovHK 2015). However, the success of introduced species management hinges upon early discovery and quick action (Simberloff et al. 2005). Research on ecological impacts of non-marine invasive species in Hong Kong has focused mainly on primary and secondary consumers such as *Pomacea canaliculata* (Fang et al. 2010;Karraker and Dudgeon 2014) and *S. invicta* (Chan and Guénard 2020), but less attention has been paid to higher level consumers such as predacious fish and decapods.

Hong Kong has several species of native freshwater decapods, but there is no native crayfish (Dudgeon 1999). However, unidentified feral crayfish have been observed in Pok Fu Lam Country Park since the early 2000s (Lee Wing Ho, pers. comm.) and again in 2016 (A. Lau, pers. observation). Feral populations of freshwater crayfish may be introduced after intentional release, unintentional introduction through escape from aquaculture, bait-bucket release, and release after being imported for educational purposes (Hobbs et al. 1989; Gherardi 2010). Crayfish are generalist omnivores and ecosystem engineers, and when introduced, generally have negative effects on recipient ecosystems. These effects can include reduction in the biomass and abundance of macrophytes, benthic invertebrates, fish, and amphibians (Twardochleb et al. 2013). Introduced crayfish species also tend to have a wider environmental tolerance than native species, which allows them to survive in extreme temperature conditions and low dissolved oxygen levels (Reynolds and Souty-Grosset 2011).

To date, no study has been conducted to confirm the identity of the reported crayfish species in Hong Kong. The objective of this study is to confirm the identity of this unidentified crayfish species and collect basic ecological information regarding this crayfish in the recipient ecosystem.

## Materials and methods

### Study site

This study was conducted in a stream pool (22°15′59.6″N, 114°08′23.2″E) and a section of a reservoir (22°15′58.8″N 114°08′21.5″E) in Pok Fu Lam Country Park in the Hong Kong Special Administrative Region, China (Fig. 1). Small stream pools with very low flow rates or nearly stagnant water are often formed along the stream length during the dry season (i.e. October to March). The stream pool sampled had a surface area of 5.17 m^2^ with a maximum depth of 0.62 m, and the reservoir’s surface area was 30565.46 m^2^. The stream pool substrate was mainly large cobbles with some boulders, while the reservoir substrate was mainly sand and clay with many boulders located along the shore. Riparian vegetation at the stream pool comprised mainly small trees and scrub; the reservoir was surrounded by tall grasses and a few shrubs. Emergent vegetation at the stream pool was dominated by *Acorus gramineus*. Average pH was 6.67 for the stream pool and 6.91 for the reservoir. Average dissolved oxygen was 8.84 and 9.22 mg/L for the stream pool and the reservoir, respectively.

**Figure. 1.**
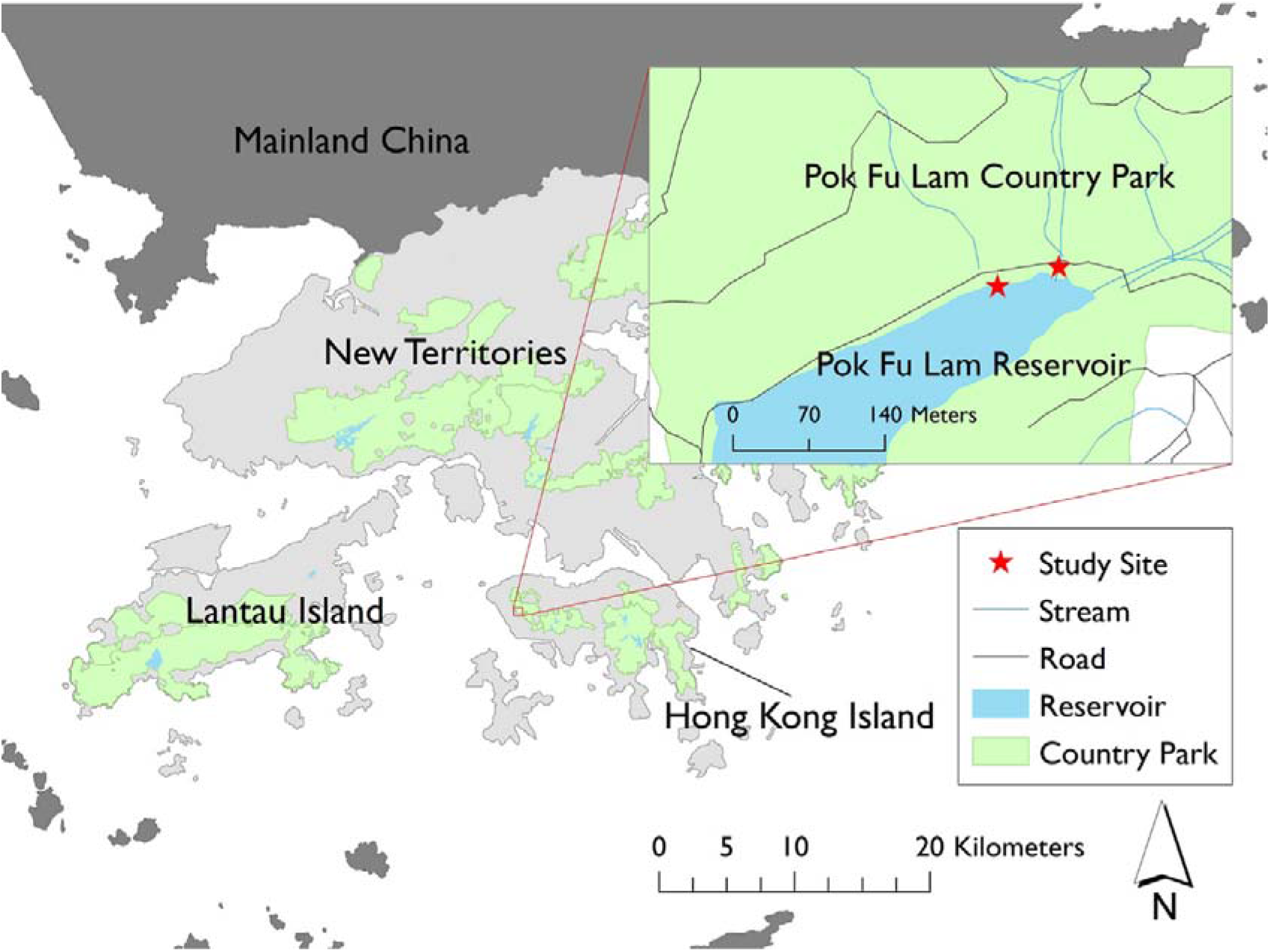
Map of Hong Kong SAR, China indicating the location of the study sites.

### Sampling

Sampling was conducted from December 2019 to February 2020. Initially, aliquots of canned cat food, minced chicken, or rice mixed with shrimp food, each weighing approximately 30 g, were placed in separate mesh bags as bait to attract crayfish. Since canned cat food appeared to be the most attractive to the crayfish (S-M Yau, pers. obs.), all subsequent sampling used only canned cat food as bait. Some crayfish were captured using hand-held dip nets when they approached bait, and some were captured with baited funnel traps (30 cm diameter, 60 cm long, 1 cm mesh size). The duration of the dip net capture period was around 6 man-hour per sampling occasion, while the funnel traps (two in the stream pool and two in the reservoir) were placed at the sampling sites for around 10 hours (overnight) per sampling occasion. In total, we surveyed two sampling sites for four occasions.

Standard morphological measurements were conducted for each crayfish captured. The total length, carapace length, and chela length of the crayfish were measured to the nearest millimetre with a vernier caliper. The mass (g) of the captured crayfish was measured with a scale. The sex of each crayfish was identified based on the position of the genital openings. The genital openings of males and females are located at the base of the fifth and third pair of pereiopods, respectively (Parnes et al. 2003). If the genital opening was located at the base of the fifth and the third pair of the walking legs, that individual was classified as intersex (Parnes et al. 2003). All crayfish captured were not released back into the sampling site, but by-catch was immediately released back into the water. The captured crayfish were transported to the laboratory using a cooler filled with ice. At the time of sampling, the water temperature was between 16.1°C and 19.1°C (mean = 18.15°C, SD = 0.86).

### Species identification

Crayfish were identified by their morphological characteristics and DNA. Muscle tissues were extracted from the chelae and walking legs of eight crayfish using sterile scissors (Scalici et al. 2009). Genomic DNA was extracted by using the DNeasy Blood and Tissue Kit (QIAGEN, Hilden, Germany). The universal primers HCO2198 (5’-TAAACTTCAGGGTGACCAAAAAATCA-3’) and LCO1490 (5’-GGTCAACAAATCATAAAGATATTGG-3’) were used to amplify the mitochondrial cytochrome c oxidase subunit I (COI) gene by conducting polymerase chain reaction (PCR) (Folmer et al. 1994). Identical COI gene fragments with approximately 650 base pairs were amplified after PCR (Scalici et al. 2009). The confirmed PCR product was sent to the laboratory company (Tech Dragon Limited) to conduct DNA sequencing. Basic Local Alignment Search Tool (BLAST) (https://blast.ncbi.nlm.nih.gov/Blast.cgi) was used to confirm the crayfish species by comparing the 624 bps obtained sequences to sequences in the BLAST GenBank nucleotide database.

## Results

A total of 49 crayfish were captured from 4 sampling occasions (Table S1). The captured crayfish were identified as Australian redclaw crayfish, *Cherax quadricarinatus* (von Martens 1868), (Fig. 2) based on morphological characteristics and DNA barcoding. All eight sequences matched with the *C. quadricarinatus* sequences in Genbank, with similarity values ranging from 99% to 100% (Genbank Accession number: KY745779.1 and KX377348.1).

**Figure 2.**
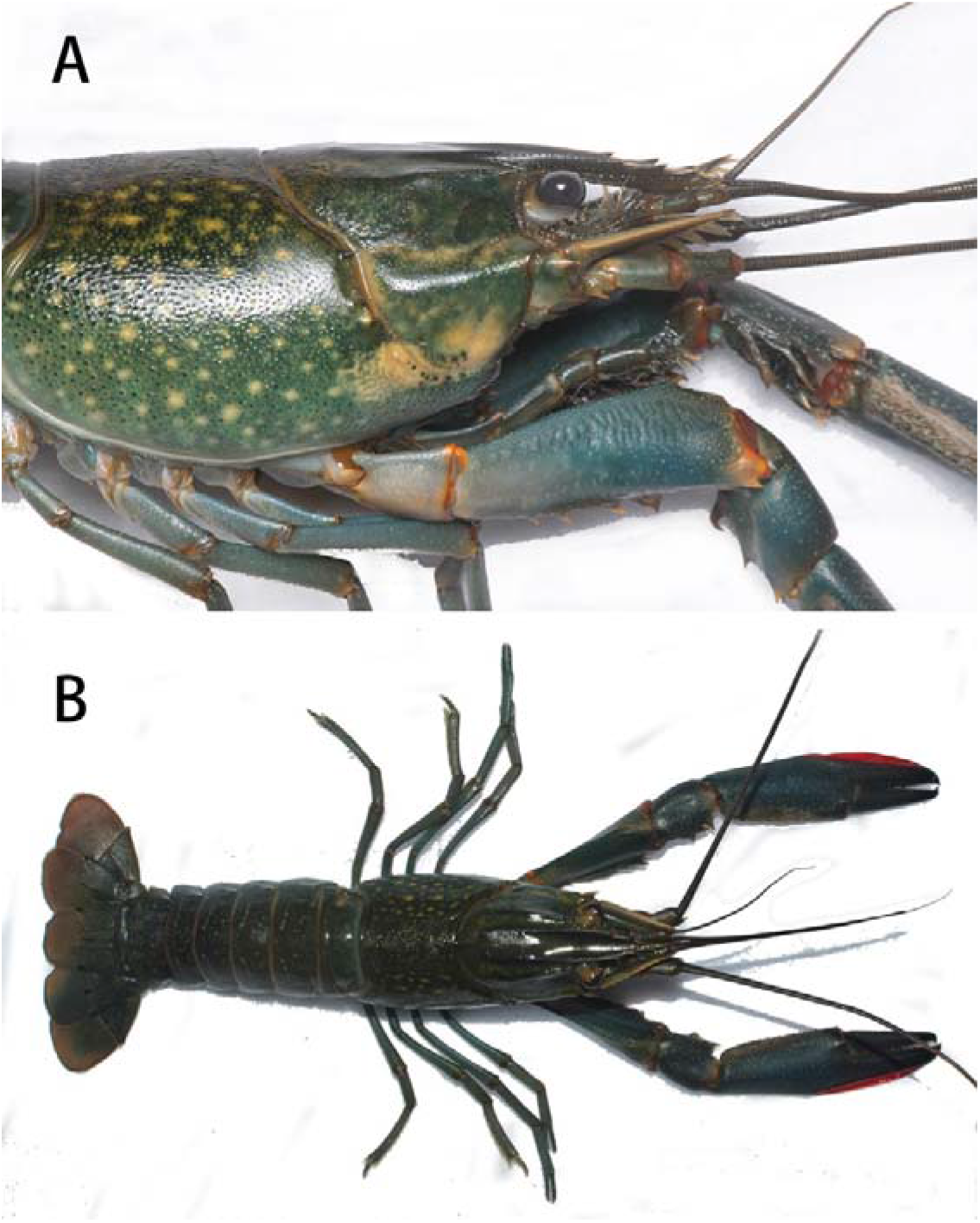
Lateral (A) and dorsal (B) views of a male *Cherax quadricarinatus* captured in Pok Fu Lam Country Park, Hong Kong SAR, China.

Total length of captured *C. quadricarinatus* was between 33 mm and 119 mm (mean = 65.06, SD = 21.28 mm). Carapace length ranged from 17.20 mm to 56.40 mm (mean = 30.70, SD = 10.13 mm), while length of chela was between 8.60 mm and 43.40 mm (mean = 20.78, SD = 9.06 mm) (Fig. 3). A total of 29 males and 19 females and 1 intersex were captured; the sex ratio was not significantly different from 1:1 (χ^*2*^ = 1.653, *p* = 0.199). We captured 33 crayfish from the stream pool and 16 crayfish from the reservoir. The capture per unit effort (CPUE) for baited traps was 0.33 and 0.50 crayfish per trap night for the stream pool and the reservoir, respectively.

**Figure 3.**
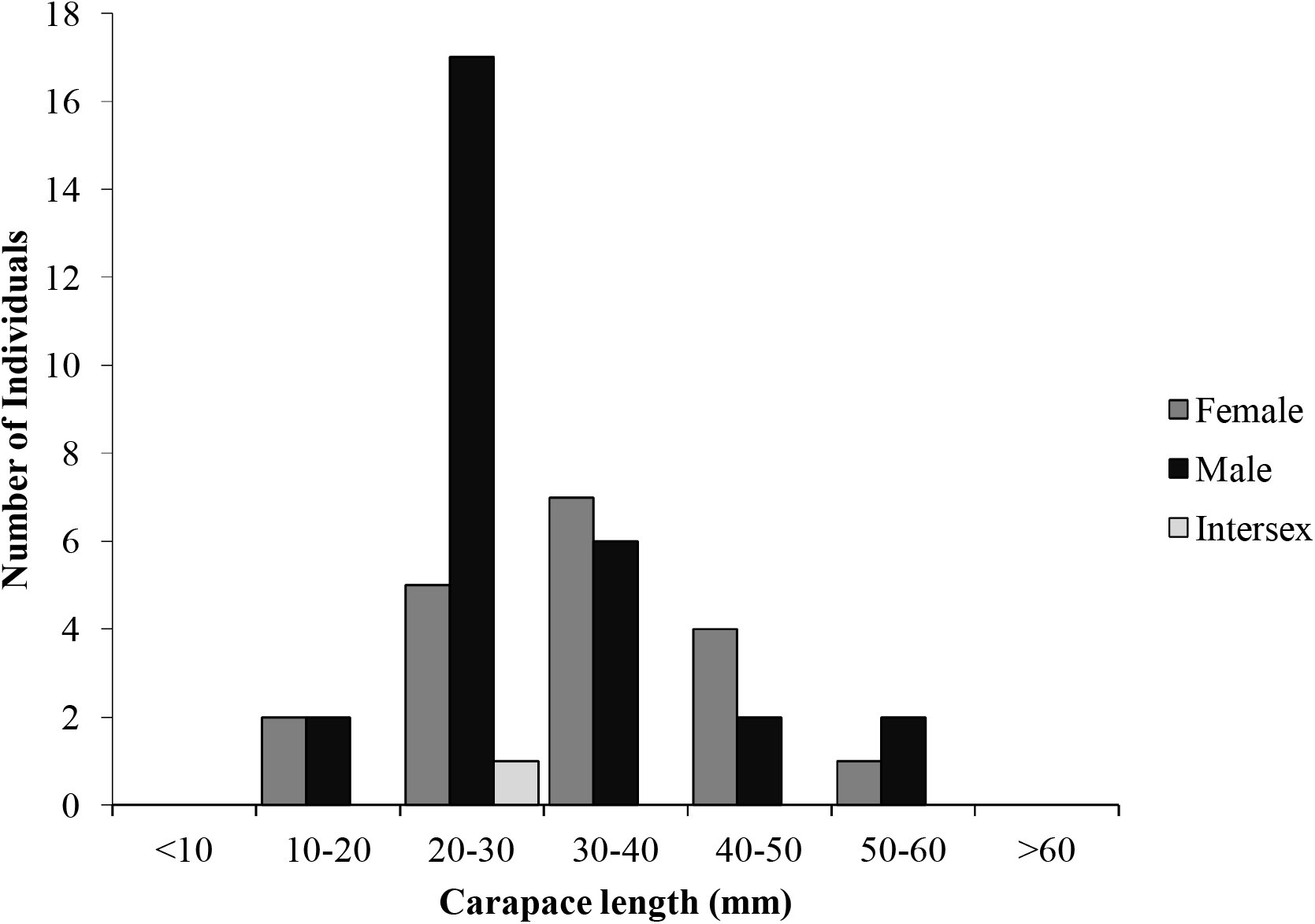
Carapace length distribution of *Cherax quadricarinatus* (n = 49) captured in Pok Fu Lam Country Park, Hong Kong SAR, China.

## Discussion

Here we provide evidence of the introduction of *Cherax quadricarinatus* in a stream and a reservoir in Pok Fu Lam Country Park in the Hong Kong Special Administrative Region, China. *Cherax quadricarinatus* is a large and conspicuously coloured crayfish native to south-eastern Papua New Guinea and northern Australia (Jones 2011). *Cherax quadricarinatus* has been introduced to many countries around the world for aquaculture, fisheries and ornamental pet trade (Ackefors 2000; Belle et al. 2011; Lodge et al. 2012; Faulkes 2015; Madzivanzira et al. 2020). Escape from captivity and deliberate release by pet owners has resulted in the establishment of feral populations globally (Lodge et al. 2012; Madzivanzira et al. 2020).

The most likely introduction pathway of *C. quadricarinatus* to Hong Kong is the deliberate release by pet owners. Live *C. quadricarinatus*, along with several other non-native crayfish species, can be found for sale in several shops in the popular “Goldfish Street” pet market in Hong Kong. Although Hong Kong imports up to 30% of all the aquacultured *C. quadricarinatus* from Australia (Jones 2011), this species is rarely consumed as food within Hong Kong. We suspect that most *C. quadricarinatus* are imported as frozen “seafood” and re-exported to China where consumption of crayfish is common. Pok Fu Lam Country Park is an apparent ‘dumping ground’ for unwanted pets, probably due to its location and ease-of-access. Over the course of this study, several other introduced species were also observed in Pok Fu Lam Country Park Reservoir and stream pools, which included: the red-eared slider *Trachemys scripta elegans*, Chinese stripe-necked turtle *Mauremys sinensis*, goldfish *Carassius auratus*, an unidentified species of jewel cichlid *Hemichromis* sp., and a breeding population of Chinese water dragons *Physignathus cocincinus*.

This study observed *C. quadricarinatus* of multiple size classes as well as a berried a female with hatchlings in July 2020, indicating that the population of *C. quadricarinatus* in Pok Fu Lam Country Park is reproducing. Most of the *C. quadricarinatus* captured were small (< 40 mm carapace length). However, this may not reflect the overall size distribution of the population. Based on our observations, most crayfish were hiding in shelters when we first arrived at the sampling sites. Individuals attracted by the bait would emerge from their shelters and thus were more likely to be captured. In contrast, the majority of *C. quadricarinatus* captured by Nunes et al. (2017) using baited crayfish traps ranged between 40 to 70 mm in carapace length. This may reflect a bias towards smaller individuals in our study. However, the South African population sampled in Nunes et al. (2017) is a long-established invasion core, and the individuals sampled are likely to be larger anyway than in recently introduced populations. Continuous monitoring is required to confirm this suspected discrepancy.

*Cherax quadricarinatus* may threaten native species through interspecific competition for food and shelter. For example, Zeng et al. (2019) demonstrated that *C. quadricarinatus* outcompetes the native freshwater crab *Parathelphusa maculata* for shelter and that this interaction leads to an overall displacement of the native species from areas with more crayfish. A study in South Africa demonstrated that biotic resistance to *C. quadricarinatus* by native, trophically analogous, freshwater crabs was size and sex-dependent. For example, the female *Potamonautes perlatus* native to Southern Africa, have significantly stronger maximum chela force than female *C. quadricarinatus* and both sexes of another introduced crayfish species *Procambarus clarkii* (South et al. 2020). In Hong Kong, *C. quadricarinatus* may compete with functionally similar native decapods such as *Macrobrachium hainanense*, *M. nipponense*, and *Nanhaipotamon hongkongense*. Unfortunately, outside of South Africam studies on the ecological impact of *C. quadricarinatus* is severely limited (Madzivanzira et al. 2020; Morningstar et al. 2020). Pinder et al. (2019) suggested that this species may deplete macrophyte cover and alter invertebrate communities in Western Australia, while Williams et al. (2001) suggested it may out compete and replace freshwater shrimp in Puerto Rico. Further studies on the potential impacts of *C. quadricarinatus* on native species are warranted.

Because of the potential negative impacts presented by *C. quadricarinatus*, it is necessary to conduct a comprehensive survey of *C. quadricarinatus* in Hong Kong in order to determine the extent of invasion to other streams and reservoirs. In South Africa and Swaziland, *C. quadricarinatus* has been shown to be capable of spreading at a rate of 8 km/year downstream and 4.7 km/year upstream (Nunes et al. 2017). Since most of the streams in Pok Fu Lam Country Park are connected through the reservoir, it is highly likely that *C. quadricarinatus* will colonize all the streams draining into the basin. In Singapore, since it was first reported in 2007, this species has spread to at least 3 of 13 reservoirs (Ahyong and Yeo 2007; Belle and Yeo 2010; Belle et al. 2011). Therefore, to detect new invasion fronts of this species, the local authority in Hong Kong should consider employing environmental DNA (eDNA) techniques, which have been demonstrated to be feasible and reliable elsewhere (Cai et al. 2017).

Targeted plans to manage this invasive species are needed to control their population size. Since pet abandonment is one likely reasons leading to the occurrence of the feral crayfish population in Hong Kong, strengthening public education is suggested to raise conservation awareness in order to solve the problem from the root. Additional legislation (with fines) that prohibits the release of animals and resources to enable enforcement of the legislation should also be considered to further discourage people from releasing animals into the wild.

## Acknowledgements

We thank Hau Ka Ki Anthony and Au Ming-Fung Franco for assistance with field work. We thank the Agriculture, Fisheries and Conservation Department for issuing a permit to collect crayfish for this study. We thank the anonymous reviewers for providing helpful comments on earlier drafts of this manuscript. We thank Ackley Lane for editing.

## Funding Declaration

The authors received no specific funding for this work.

## Authors’ Contribution

SY: research conceptualization; research conceptualization; investigation and data collection; data analysis and interpretation; writing - original draft.

AL: research conceptualization; research conceptualization; investigation and data collection; data analysis and interpretation; writing - review & editing.

## Declaration of Interests

None.

## Ethics and Permits

Ethics was not required. The Agriculture, Fisheries and Conservation Department granted permission to possess funnel traps for ecological surveys within Pok Fu Lam Country Park.

## Notes

### Competing Interest Statement

The authors have declared no competing interest.

